# Targeted degradation of SARS-CoV-2 via the autophagy-lysosome system using chemical mimetics of the N-degron pathway

**DOI:** 10.1101/2025.09.01.673409

**Authors:** Gee Eun Lee, Tae Hyun Bae, Uni Park, Jihoon Lee, Dong Won Lee, Young Jo Song, Se Hun Gu, Chi Ho Yu, Minsoo Kim, Luis Martinez-Sobrido, Murim Choi, Jun-Gyu Park, Nam-Hyuk Cho, Yong Tae Kwon, Ki Woon Sung

**Author notes:** Corresponding authors Yong Tae Kwon. Department of Biomedical Sciences, College of Medicine, Seoul National University, Seoul 03087, Korea. <u>Telephone</u>: 82-2-740-8547. <u>Fax</u>: 82-2-3673-2167. <u>Email</u>. Ki Woon Sung. Duke Center for Neurodegeneration and Neurotherapeutic, Duke University, Durham, NC, 27710, USA. <u>Telephone</u>:1-684-5224. <u>Email</u>. Nam-Hyuk Cho. Department of Microbiology and Immunology, College of Medicine, Seoul National University, Seoul 03087, Korea. <u>Telephone</u>: 82-2-740-8392. <u>Fax</u>: 82-2-743-0881. <u>Email</u>. Jun-Gyu Park, Laboratory of Veterinary Zoonotic Diseases, College of Veterinary Medicine, Chonnam National University, Yongbong-ro 77, Buk-gu, Gwangju 61186, Korea. <u>Telephone</u>: 82-62-530-2877. <u>Email</u>. These authors equally contributed to this work (Gee Eun Lee, Tae Hyun Bae, Uni Park).

## Abstract

In the N-degron pathway, ATE1 transfers the amino acid L-arginine (L-Arg) from Arg-tRNA^Arg^ to N-terminal (Nt) residues of cellular proteins. The resulting Arg/N-degrons bind the autophagic receptor p62/SQSTSM-1/Sequestosome-1 to induce lysosomal degradation of various biomaterials. Here, we demonstrate that the chemical mimetics of Arg/N-degrons, termed autophagy-targeting ligands (ATLs), can induce lysosomal degradation of SARS-CoV-2 (severe acute respiratory syndrome coronavirus-2) via p62-mediated macroautophagy. In Vero E6 cells infected with SARS-CoV-2, ATLs promoted p62 self-polymerization and enhanced LC3 synthesis and lipidation, leading to viral sequestration within autophagosomes for lysosomal degradation. In transgenic mice overexpressing human angiotensin-converting enzyme 2 (ACE2), oral administration of ATL1014 inhibited viral replication and increased viability. In a Syrian hamster model, ATL1014 attenuated viral replication in the lungs and demonstrated efficacy in inflammatory lesions and pulmonary congestions. These results identify the N-degron pathway as a potential target for a host-targeting strategy (HTS) against a broad spectrum of viruses.

## Introduction

COVID-19 caused more than 700 million cases with approximately 7.1 million deaths globally by 2025, making it the deadliest pandemic in recent history.^1^ The disease, caused by SARS-CoV-2, is characterized by pneumonia-associated symptoms.^2^ Belonging to the family of *Coronaviridae*, SARS-CoV-2 infects alveolar epithelial cells through ACE2 and requires spike (S) protein priming mediated by the cellular serine protease TMPRSS2.^3^ Upon entry, the virus hijacks host cell membranes to form double-membrane vesicles (DMVs).^4^ Virus-induced DMVs provide a protective environment for viral RNA intermediates during replication and shield them from recognition by innate immune sensors in the host cell. Rapid proliferation of viruses triggers dysregulated immune responses and hyperactivation of pro-inflammatory cytokines, ultimately leading to cytokine storm.^5,6^ This excessive inflammation contributes to acute respiratory distress syndrome (ARDS), multi-organ failure, and pathological features such as pulmonary edema, fibrosis, and impaired gas exchange.^7,8^ Within a couple of years since the pandemic arose, several variants of concern (VOC) have emerged, including Alpha (B.1.1.7), Beta (B.1.351), Delta (B.1.617.2), Gamma (P.1), and Omicron (B.1.1.529).^9^ Although effective vaccines for COVID-19 have been developed in unprecedented periods of time as a preventative tool in battling against COVID-19,^10^ the rise of variants has restricted their effectiveness and allowed evasion.^11^ Additionally, concerns have arisen regarding mRNA vaccines, including the shedding of spike proteins or its peptide fragments into the circulation, which may interact with host proteins or other cellular targets.^12^ These challenges underscore the need for strategies that can overcome vaccine limitations and effectively address future variants, as currently available vaccines may be insufficient against newly emerging strains.

Central to infectious diseases is the capacity of viruses to evade host defense systems, enabling them to propagate to other cells in variable forms.^13^ Traditional approaches in antiviral drug discovery focus on small molecules that bind specific viral proteins but non-existing in host proteins.^14^ Although extensive efforts have been made to develop antiviral drugs, successful treatments have been limited to a handful of viruses, such as human immunodeficiency virus (HIV), hepatitis C virus, and herpes simplex virus.^15^ Remdesivir, an antiviral nucleoside analog, binds to the viral RNA-dependent RNA polymerase (RdRp) and inhibits the replication of SARS-CoV-2.^16^ Despite extensive effort, the major difficulty in antiviral development is the rapid emergence of drug-resistant genetic variants, which hinders the long-term efficacy of direct-acting antivirals (DAAs).^17^ To cope this challenge, HTS offers a robust alternative by targeting conserved host factors critical for viral replication.^18,19^ Unlike DAAs, HTS confers broad-spectrum antiviral efficacy, reduces the likelihood of resistance due to viral mutations, and provides potential solutions for emerging variants.^20^ For example, a TMPRSS2 inhibitor has shown potential broad-spectrum applications against respiratory viruses.^21–23^ However, most HTS studies remain largely at a proof-of-concept level, requiring further studies to advance into clinical applications.

The N-degron pathway mediates the degradation of various biomaterials via the ubiquitin ^24^-proteasome system (UPS) or the autophagy-lysosome system (ALS).^25–27^ Central to the pathway is Nt-arginylation, in which ATE1 R-transferase (EC 2.3.2) transfers the L-Arg from Arg-tRNA^Arg^ to Nt-residues, such as aspartate (Asp), glutamate (Glu), and oxidized cysteine (Cys).^28^ The resulting Arg/N-degrons are recognized and bound by specific N-recognins that target biomaterials to the UPS or ALS.^29,30^ In the UPS, Arg/N-degrons are recognized by cognate N-recognins that promote substrate ubiquitination and proteasomal degradation.^31^ We have shown that Arg/N-degrons also modulate lysosomal destruction through their recognition by the N-recognin p62 that functions as an autophagy receptor.^32^ Upon binding to the ZZ domain of p62, Arg/N-degrons induce a conformational change to facilitate self-polymerization of p62 in the complex with its cargoes and p62 interaction with LC3 on autophagic membranes.^29^ In parallel, a subpopulation of Nt-Arg-bound p62 induces omegasome biogenesis to receive incoming autophagic cargoes.^29^ Based on this mode-of-action, we also developed chemical mimetics of Arg/N-degrons as ATLs that bind p62 via N-degron interactions.^29,33^ ATLs were demonstrated to facilitate lysosomal degradation of misfolded protein aggregates,^34–36^ the ER,^37^ the mitochondrion,^38^ intracellular bacteria,^39^ and lipid droplets.^40^

This study was designed to explore the N-degron pathway as a HTS to degrade intracellular viruses via p62-dependent autophagy. We demonstrate that chemical mimetics of Arg/N-degrons activate p62, which in turn targets SARS-Cov2 to autophagic membranes leading to their lysosomal co-degradation. In a mouse model of SARS-CoV-2 infection, ATL1014 inhibited viral infectivity and reduced inflammatory damages in the lungs, leading to increased viability. In Syrian hamsters infected with SARS-CoV-2, ATL1014 mitigated viral infectivity and alleviated lung inflammation. Our results identify the N-degron pathway as a potential HTS target to develop a broad-spectrum antiviral approach.

## Results

### Chemical mimetics of Arg/N-degron targets SARS-CoV-2 via p62-dependent autophagy

To determine the impact of SARS-CoV-2 infection on the ALS, we monitored the autophagic flux in Vero E6 epithelial cells derived from the kidney of an African green monkey.

Immunoblotting showed an unusually increased level of p62 upon viral infection (Figure S1A). When visualized using immunofluorescence staining, the cells excessively accumulated p62^+^ puncta in the cytosol, which failed to enter autophagic flux into LC3^+^ phagophores (Figure S1B). These results suggested that the degradative flux in p62-mediated ALS was suppressed upon SARS-CoV-2 infection (Figure S1A-S1D). We therefore determined whether intracellular SARS-CoV-2 can be directly targeted for lysosomal degradation through the chemical activation of p62 as an N-recognin of the Arg/N-degron pathway. Screening of ATL derivatives to elevate autophagic activities in the infected Vero E6 cells identified two previously developed ATLs (ATL1024, and ATL1038) for further characterization (Figure 1A).^39^ We also modified these ATLs to design a new compound, ATL1014, that can be orally administered in animals (Figure 1A).

**Figure 1.**
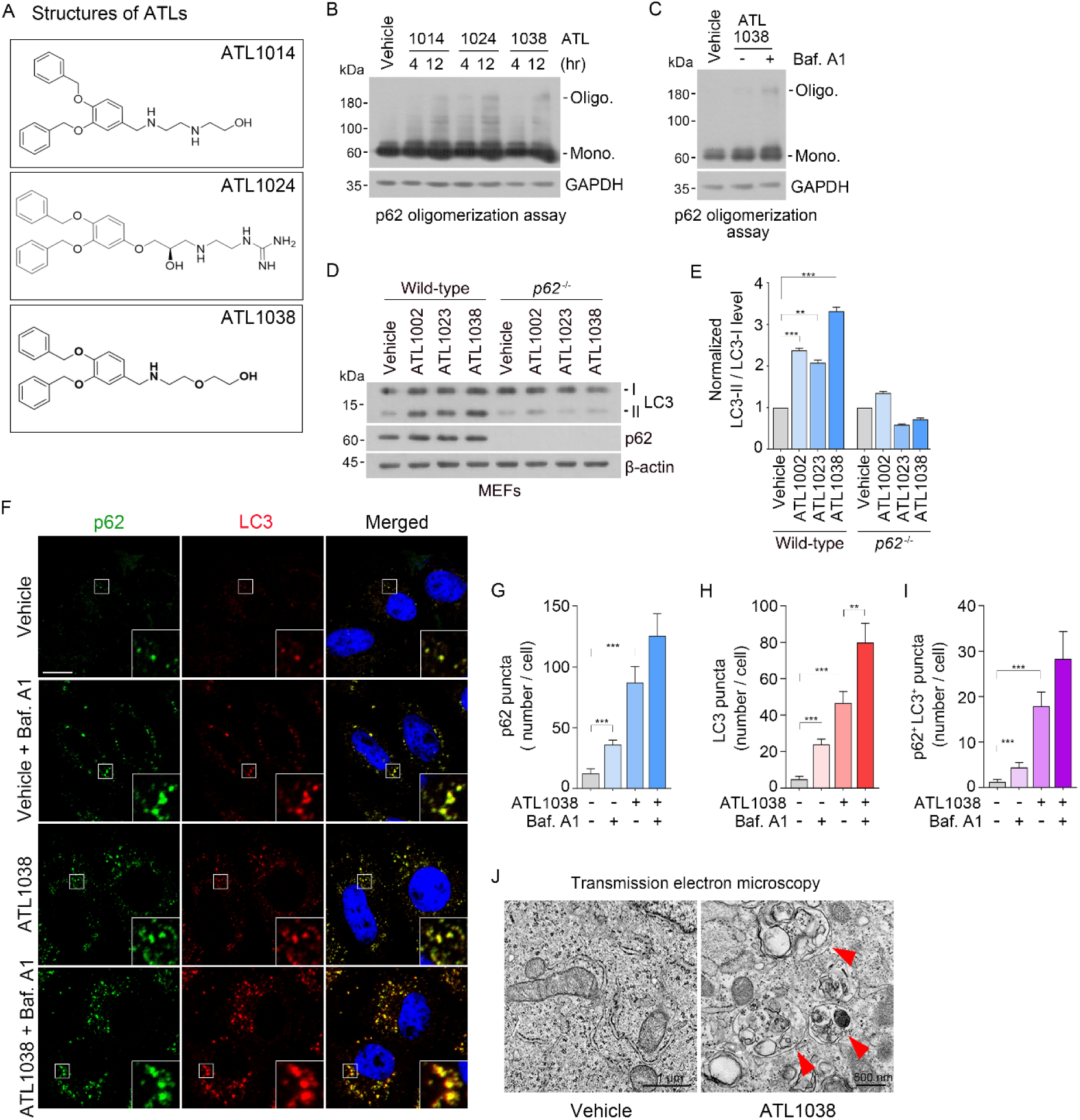
Development of antiviral chemicals that exert efficacy via p62-depedent autophagy. (A) The chemical structures of ATL1014, ATL1024, and ATL1038. (B-C) *In vivo* p62 oligomerization assay in HEK293T cells (B) treated with ATL1014, ATL1024, and ATL1038 (all 1 μM) and (C) treated with ATL1038 (1 μM, 4 h), with or without bafilomycin A1 (200 nM, 4 h). (D) WT and p62^-/-^ MEF cells treated with DMSO (vehicle), ATL1002, ATL1023 as a positive control and ATL1038 (all 1 μM, 4 h). (E) Quantification of (D). (F) Immunofluorescence images of p62 and LC3 colocalization in Vero E6 cells treated with ATL1038 (1 μM, 12 h), with or without bafilomycin A1 (200 nM, 4 h). Scale bar, 10 μm. (G-I) Quantification of p62 puncta (G), LC3 puncta (H), and p62^+^LC3^+^ puncta (I) in each cell. (J) TEM images of Vero E6 cells treated with ATL1038 (1 μM, 24 h). Red arrowheads indicate autophagosome. Scale bar, 500 nm. For all quantifications, error bars represent SEM. *, p < 0.05; **, p < 0.01; ***, p < 0.001.

To compare the activity of ATLs as p62 agonists, we measured the *in vivo* self-oligomerization of p62. HEK 293T cells treated with ATLs were analyzed using non-reducing SDS-PAGE to separate oligomeric and high molecular species of p62 from its monomers.^32,34,41^ Immunoblotting analyses showed that ATLs promoted the self-polymerization of p62, highlighting its role as a critical marker of p62 activity (Figure 1B). High-molecular species of p62 further accumulated when autophagic flux was blocked with bafilomycin A1, an inhibitor of the lysosomal V-ATPase (Figure 1C).^42^ These results collectively support that ATL1038 directly induces engagement with p62. Importantly, the ability to induce LC3-II formation was abolished in *p62^-/-^* mouse embryonic fibroblasts (MEFs) (Figures 1D and 1E). The immunofluorescence staining analysis provided further confirmation of ATL1038’s ability to stimulate the formation of p62^+^ and LC3^+^ cytosolic puncta (Figures 1F-1I). The accumulation of p62^+^LC3^+^ cargoes increased following autophagy inhibition by bafilomycin A1 (Figures 1F and 1I). When visualized using transmission electron microscopy, the activation of LC3 correlated to the biogenesis of LC3^+^autophagic membranes (Figure 1J). These results demonstrate that ATLs activate p62-dependent autophagy in Vero E6 cells infected with SARS-CoV-2.

In the docking simulation study, ATLs all located in the position to which the first and second residues of the N-degron substrates bind (Figures S2A-S2D). ATL1014 and ATL1038 exhibited a binding mode that partially mimicked the interaction of p62 ZZ domain with its physiological N-degrons (Figures S2B and S2D). Both compounds formed hydrogen bonds between their two nitrogen atoms and the side chains of Asp129 and Asp149, respectively, with a T-shaped π-π interaction was observed between Phe168 and the phenyl ring of both ATL1014 and ATL1038. Similar to ATL1014, the hydroxyl moiety of ATL1024 formed a hydrogen bond to the side chain of Asp147, and its guanidinium moiety formed a salt bridge with the side chain of Asp129 (Figure S2C).

To assess the antiviral activity of ATLs, Vero E6 cells infected with SARS-CoV-2 (0.01 MOI) were subjected to immunofluorescence staining analyses, in which the levels and distributions of nucleocapsid (N) protein correlated into dose-response curves using an in-house software.^43^ ATL1014 and ATL1038 exhibited significant antiviral efficacy with the half maximal inhibitory concentration (IC_50_) values of 5.7 μM and 11.7 μM, respectively (Figures 2A and 2C). In sharp contrast, significant efficacy was not observed with ATL1024 (Figure 2B). To compare the antiviral efficacy and cytotoxicity of ATLs, we measured the mean 50% cytotoxicity concentration (CC_50_) values, which showed 14.1 μM and 28.4 μM for ATL1014 and ATL1038, respectively (Figure S3A and S3C). As an alternative approach, we also performed plaque-reduction assays with cells infected with SARS-CoV-2 at 0.01 MOI for 2 h. Following the treatment of ATLs for 3 days, plaques were visualized and quantified after staining for N protein. ATL1014 showed potent antiviral activity, with 92% inhibition observed at 10 μM (Figures 2D and 2E). Similarly, ATL1038 revealed marked reduction in the number of N protein^+^ plaques at 10 μM and 20 μM, with 26% and 76% inhibition, respectively (Figures 2F and 2G). These results demonstrate the antiviral efficacy of ATL1014 and ATL1038 against SARS-CoV-2.

**Figure 2.**
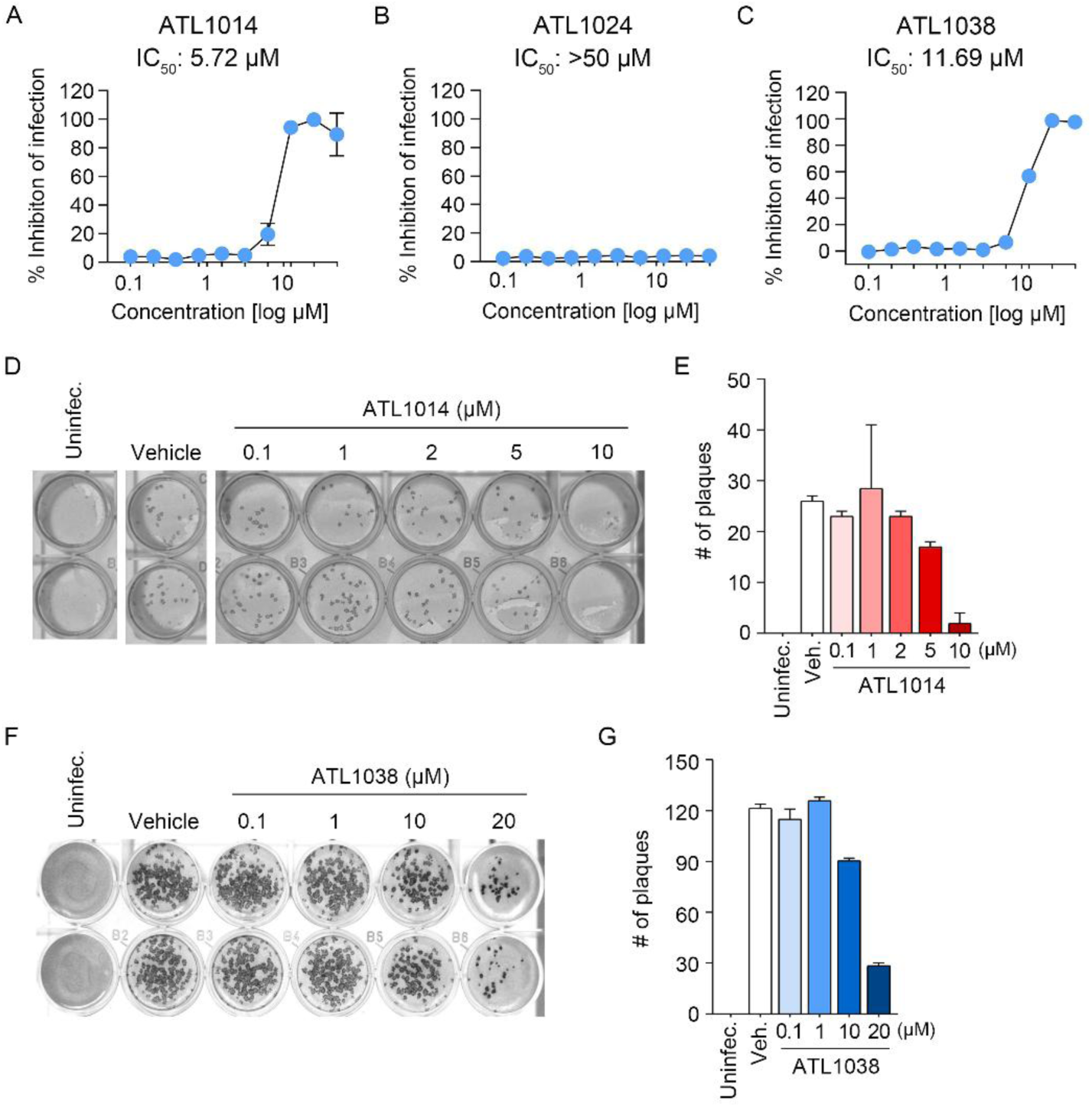
ATLs inhibit infectivity of SARS-CoV-2. (A-C) Dose-response curve analysis for antiviral efficacy of ATL1014, ATL1024, and ATL1038. (D and F) Vero E6 cells infected with SARS-CoV-2 were incubated with ATL1014 (D) and ATL1038 (F) at the indicated concentrations for 70 hours. The number of plaques was counted using N protein antibody. (E and G) Quantification of plaque numbers for ATL1014 (E) and ATL1038 (G).

Next, we compared the activities of ATLs with antiviral drugs available in the market. Remdesivir showed IC_50_ of 7.2 μM. Chloroquine (9.7 μM) and Lopinavir (11.5 μM) also inhibited viral replication with similar efficacy (Figures S4A-S4C). These results suggest that the antiviral activities of ATLs *in vitro* are largely comparable to antiviral drugs in the market.

### Chemical Arg/N-degrons of p62 induce the lysosomal degradation of SARS-CoV-2

Upon SARS-CoV-2 infection, viral RNAs are replicated within ER-derived DMVs.^44^ Following replication, viral components self-assemble with structural proteins at the ERGIC (ER-Golgi intermediate compartment) or the Golgi apparatus. Following budding off, the newly formed viral particles are trafficked in single membrane vesicles (SMVs) and eventually released from the host cell via the exocytic pathway.^45,46^ To determine whether ATL1038 induces autophagic degradation of SARS-CoV-2, we visualized intracellular SARS-CoV-2 in Vero E6 cells using TEM analyses. Following infection, the number of viral DMVs with diameters of 200-300 nm increased predominantly in the perinuclear region (Figure 3A). Importantly, ATL1038 treatment significantly reduced the number of DMVs that support viral replication (Figures 3A and 3B). Substantial numbers of assembled viral particles were observed in the extracellular space due to exocytosis following SARS-CoV-2 infection (Figure 3C). In ATL1038-treated cells, a reduction of viral particles and those encapsulated within SMVs were observed. Consistently, further analysis revealed that surviving viruses were typically sequestered by autophagic membranes, a portion of which were in the process of lysosomal fusion (Figure 3C). When autophagic flux was blocked with bafilomycin A1, viral particles massively accumulated within the lumen of autophagosomes and autolysosomes (Figure 3C). These results suggest that ATL1038 redirects SARS-CoV-2 from its normal life cycle to autophagic flux, facilitating the targeting of SARS-CoV-2 to autophagic membranes for lysosomal degradation.

**Figure 3.**
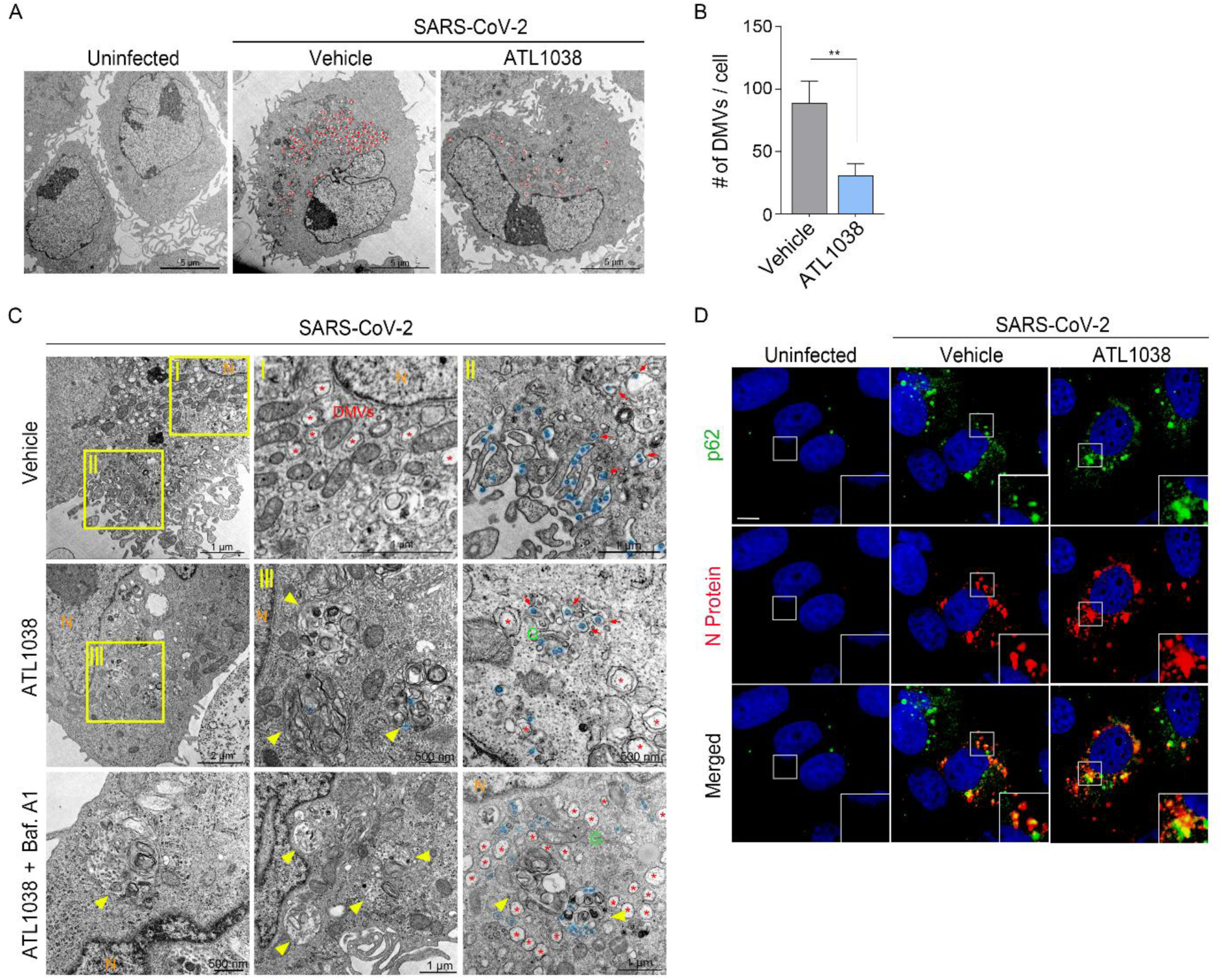
Chemical Arg/N-degrons of p62 induce the lysosomal degradation of SARS-CoV-2. (A) TEM images of Vero E6 cells infected with SARS-CoV-2 for 2 h, followed by treatment with or without ATL1038 (10 μM, 12 h). Red asterisks indicate DMVs. (B) Quantification of DMVs. (C) TEM images of infected Vero E6 cells treated with or without ATL1038 (10 μM, 12h), and with or without bafilomycin A1 (200 nM, 4 h). Red asterisks indicate DMVs and red arrows mark viral particles within SMVs. Yellow arrowheads point to autophagic vesicles. Viral particles are colorized blue. Letter N in orange represents nucleus and letter G in green represents golgi apparatus. (D) Immunofluorescence images of p62 and N protein colocalization in Vero E6 cells uninfected or infected with SARS-CoV-2 were treated with or without ATL1038 for 12 hours under confocal microscopy. Scale bar, 10 μm. For all quantifications, error bars represent SEM. *, p < 0.05; **, p < 0.01; ***, p < 0.001.

As an alternative approach, N protein of SARS-CoV-2 was monitored using immunostaining analysis. N protein^+^ signals were visualized as cytosolic puncta, representing virus-containing vesicles (Figure 3D). Importantly, ATL treatments showed a reduction in both the intensity and number of N protein^+^ signals. Surviving N protein^+^ signals in ATL-treated cells were typically localized within p62^+^ autophagic membranes (Figure 3D). These results suggest that the ATL1038 promotes the lysosomal degradation of intracellular SARS-CoV-2 via p62-dependent autophagy.

### *In vivo* efficacy of ATL1014 in viral infectivity, lung damages, and viability of ACE2 transgenic mice infected with SARS-CoV-2

To test the *in vivo* efficacy of ATLs in animal models, we compared their pharmacokinetic (PK) profiles in mice. When oral administered at 10 mg/kg, the AUC_last_ values of plasma were 388.4 ± 200.9 and 39.5 ± 2.6 and h*ng/mL, respectively, for ATL1014 and ATL1038 (Table 1). In addition, ATL1014 was superior to ATL1038 in the peak concentration (60.7 ± 20.8 vs. 20.7 ± 3.6 ng/mL) and the mean plasma half-life (8.74 ± 2.33 vs. 2.33± 0.49 h) (Table 1). We therefore chose ATL1014 for animal experiments.

**Table 1.** PK parameters of ATL1014 and ATL1038.

|  | ATL1014 |  |  | ATL1038 |
| --- | --- | --- | --- | --- |
| Route | IP | PO | IV | PO |
| Dose/ Unit (mg/kg) | 10 | 10 | 1 | 5 |
| $T_{1/2}$ (h) | 13.2 | 8.7 | 6.1 | 2.3 |
| $T_{max}$ (h) | 0.08 | 0.5 | 0.08 | 0.33 |
| $C_{max}$ (ng/ml) | 757.0 | 60.7 | 223.4 | 20.7 |
| $AUC_{last}$ (h*ng/ml) | 1661.9 | 388.4 | 190.4 | 39.5 |
| F (%) | 87.3 | 20.4 |  |  |

To assess the antiviral efficacy of ATL1014 in an acute model of SARS-CoV-2 pathogenesis, we used transgenic mice, in which the keratin 18 promoter overexpresses human ACE2, a receptor for Spike of SARS-CoV-2.^47^ Five-week-old hACE2 females were intranasally infected with 10^4^ PFU (plaque forming unit) recombinant (r) SARS-CoV-2 that was genetically engineered to contain two reporters, fluorescent mCherry and the Nano luciferase (Nluc) in the nucleocapsid.^47^ The mice infected with rSARS-CoV-2 were orally administered either a vehicle or ATL1014 at 10 or 30 mg/kg, starting from a day before infection, followed by daily treatment for five days (Figure 4A). Body weight was monitored throughout the infection period. While vehicle-treated mice underwent weight loss by approximately 25% by 6 days post injection (dpi), the ATL1014 group lost weights 15% less as compared with the vehicle group (Figure 4B). Notably, 30 mg/kg ATL1014 group exhibited remarkable recovery in body weight after the initial decline, regaining weight as the infection progressed indicating protective effects on infected mice (Figure 4B). Consistent with other studies,^47–50^ all the infected mice treated with the vehicle died by 7 dpi (Figure 4C). By sharp contrast, oral administration of 30 mg/kg of ATL1014 significantly improved survival rate. Only one mouse succumbed to the infection on 4 dpi, with the remaining mice survived until 14 dpi, resulting in a final survival rate of 75% (Figure 4C). These results demonstrate that ATL1014 exerts antiviral efficacy against rSARS-CoV-2 in K18 hACE2 mice.

**Figure 4.**
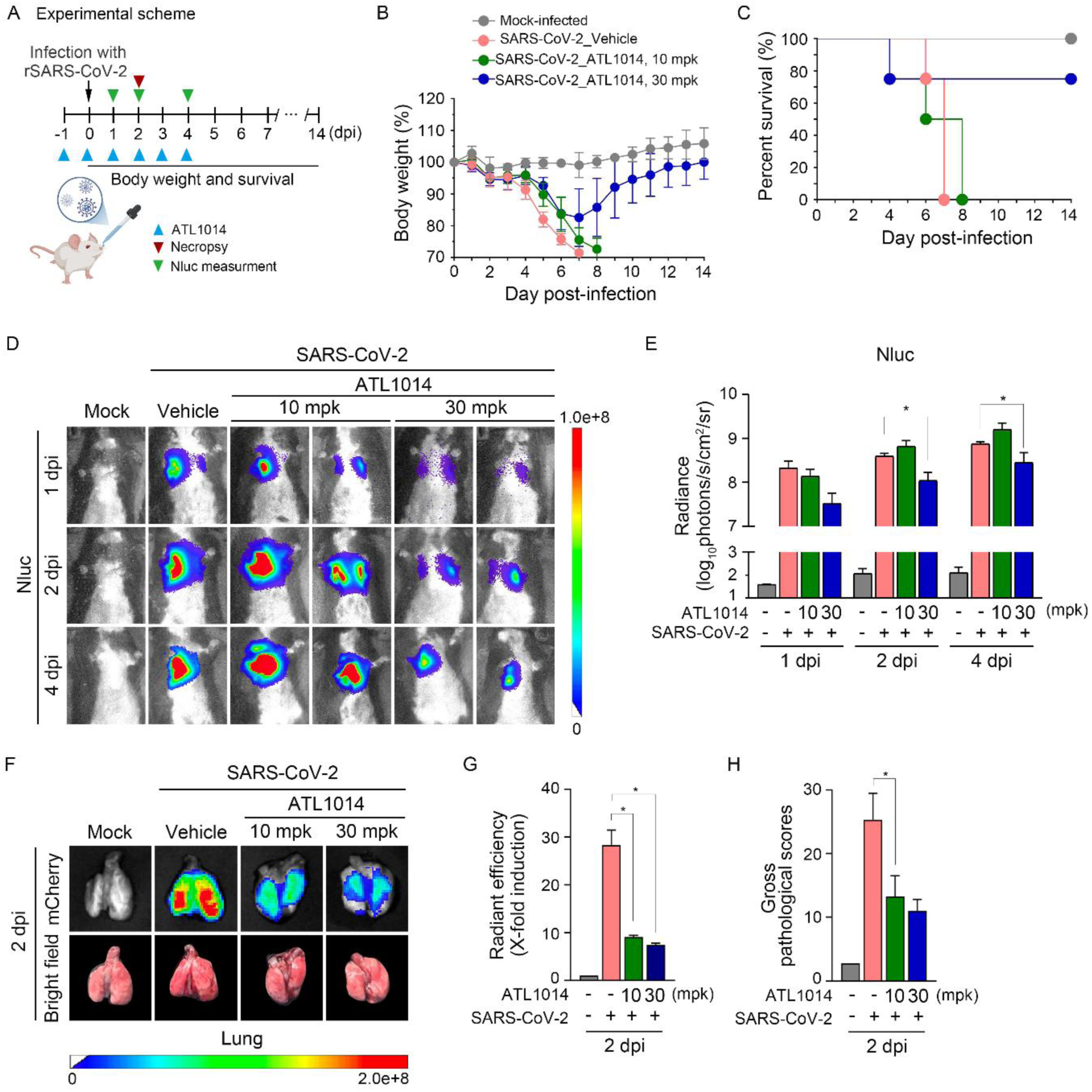
*In vivo* efficacy of ATL1014 in viral infectivity, lung damages, and viability of ACE2 transgenic mice infected with SARS-CoV-2. (A) Schematic of hACE2 transgenic mice and treatment schedule with PBS (vehicle) or ATL1014 (10 mpk and 30 mpk) orally. Mice were infected with mock or rSARS-CoV-2/WT mCherry/nLuc-2A (1 x 10^4^ PFU / mouse). (B and C) Body weight (B) and survival (C) were evaluated at the indicated DPI. (D and E) *In vivo* luminescence images of mice determined using an IVIS system at the indicated DPI. Mice were anesthetized at 1,2, and 4 dpi and retro-orbitally injected with the Nluc substrate. Nluc expression was determined using IVIS system (D) and quantitatively analyzed by the Aura program (E). (F and G) *Ex vivo* images of lungs and mCherry expression determined using IVIS system and mean values were normalized to the autofluorescence in mock-infected mice at each time point and were used to calculate fold induction by the Aura program (G). (H) Gross lesions on the lung surfaces were quantitatively analyzed by image J. For all quantifications, error bars represent SEM. *, p < 0.05; **, p < 0.01; ***, p < 0.001.

To monitor the replication of SARS-CoV-2 in mice, we measured the expression of Nluc at 1, 2, and 4 dpi using a bioluminescence imaging system (IVIS Spectrum), which captures and quantifies the luminescent signals emitted by Nluc in live animals.^51^ This approach enables real-time tracking of viral replication and dissemination across different tissues. In vehicle-treated mice, the luminescent signal from Nluc was readily detected from 1 dpi and spatiotemporally increased by 4 dpi (Figures 4D and 4E). The Nluc signals were significantly reduced by oral administration of 30 mg/kg ATL1014 throughout 4 dpi, indicating decreased viral replication (Figure 4D and 4E). The results from the Nano luciferase were reproduced when mCherry expression was monitored in the lungs harvested at 2 dpi (Figures 4F and 4G). Consistent with the reduced number of viruses, lung pathology scores were significantly lower in mice treated with 30 mg/kg ATL1014 compared with vehicle mice (Figure 4H). Collectively, these results demonstrate that oral administration of ATL1014 efficiently inhibits the replication of SARS-CoV-2 in the lungs of hACE2 mice.

### ATL1014 exhibits therapeutic efficacy in viral infectivity and lung damages of Syrian hamsters infected with SARS-CoV-2

As an alternative to validate the antiviral efficacy of ATL1014 *in vivo*, we also employed a Syrian hamster model, which has been widely utilized in SARS-CoV-2 research due to its susceptibility to infection and ability to replicate key pathological and clinical features of human COVID-19.^52,53^ This model is known to develop both upper and lower respiratory tract involvement, accompanied by histopathological changes such as alveolar inflammation and vascular congestion.^53^

The hamsters (n = 5 per group) were infected intranasally with 5 x 10^2^ PFU of SARS-CoV-2 (BetaCoV/Korea/KCDC03/2020). To evaluate the antiviral efficacy of ATL1014, the hamsters received 20 mg/kg of ATL1014 intraperitoneally two hours post-infection, followed by daily administrations for 4 consecutive days, and were sacrificed on the day 5 post-infection (Figure 5A). After 5 days of treatment, no significant differences were observed in the gross phenotypes. Subsequently, the lungs were harvested for histopathological analysis to examine the progression of pathological changes. Consistent with previous reports,^52^ the vehicle group exhibited inflammatory lesions and other morphological abnormalities, as evidenced by multifocal and locally extensive congestions across 40-50% of the lung surface (Figure 5B). Importantly, these pathological phenotypes were significantly alleviated in ATL1014-treated hamsters (Figure 5B). To further characterize the pathological symptoms, the cross sections of the lungs were histologically examined using hematoxylin and eosin (H&E) staining. Infected lungs from vehicle-treated hamsters revealed focal inflammation in the alveolar septa, characterized by infiltration of mononuclear cells, septal thickening, and congestion (Figures 5C and 5D). These inflammatory damages were associated with alveolar collapse and pulmonary consolidation. Importantly, ATL1014 exhibited robust efficacy in alleviating all these pathological symptoms (Figures 5C-5E). These results demonstrate that ATL1014 inhibits the infectivity of SARS-CoV-2 in the lung of hamsters.

**Figure 5.**
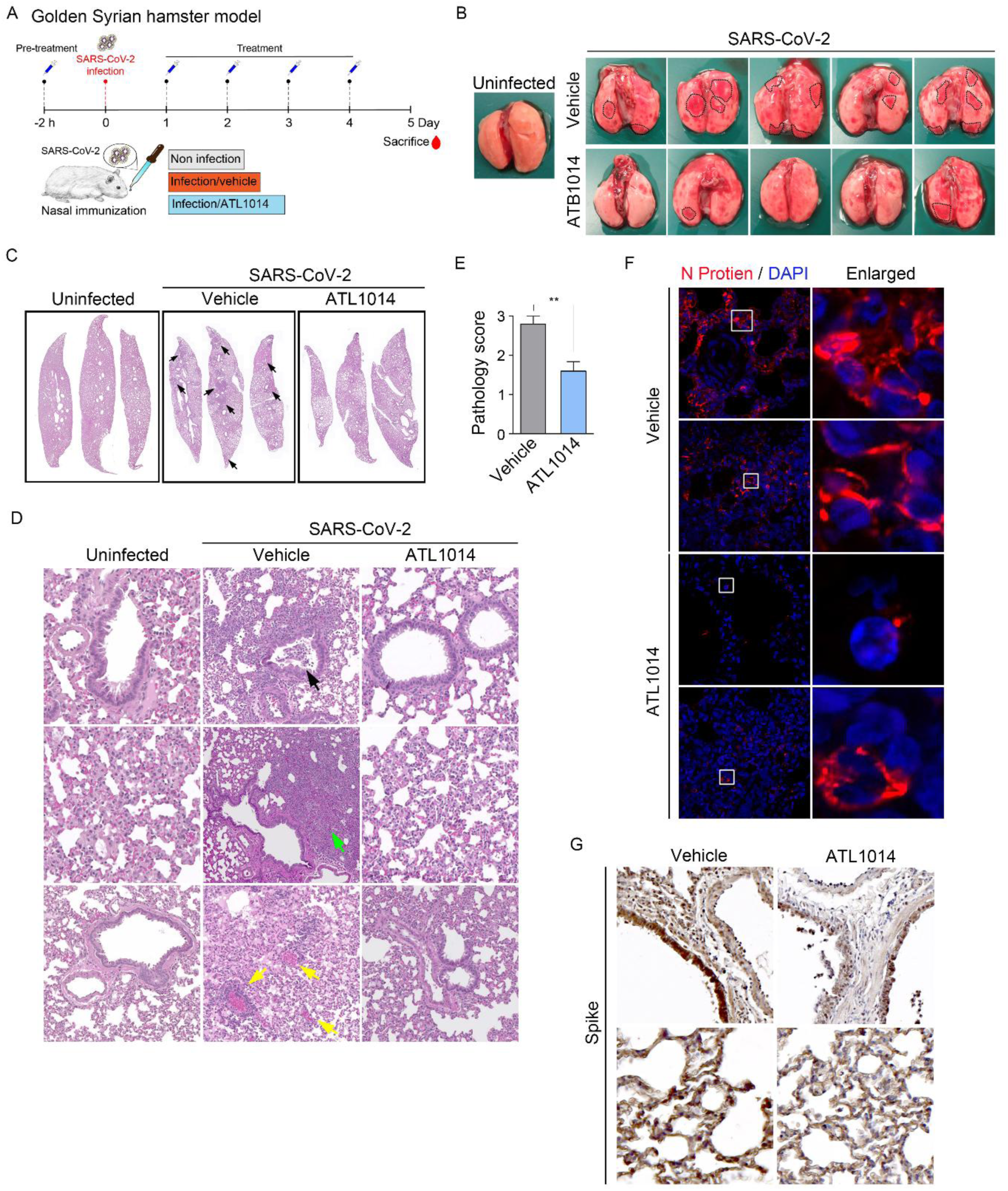
ATL1014 exhibits therapeutic efficacy in viral infectivity and lung damages of Syrian hamsters infected with SARS-CoV-2. (A) Schematic of hamster experiment. (B) Pictures of the lungs of Golden Syrian hamsters infected with mock or SARS-CoV-2 (5 x 10^2^ PFU) followed by intraperitoneally treatment of vehicle or ATB1014 (20 mg / kg) 5 times. Black dotted lines indicate lung lesions. (C) Hematoxylin and eosin analysis of the lung. Black arrows indicate histopathological lesions. (D) Histopathological analysis of lungs. Black arrow indicates cell debris. Green arrow indicates congestion. Yellow arrows indicate hemorrhage. (E) Graph represents the pathological score. (F) Immunofluorescence analysis for N protein of SARS-CoV-2 in the lungs. (G) Immunohistochemical analysis for S protein of SARS-CoV-2 in the lungs. For all quantifications, error bars represent SEM. *, p < 0.05; **, p < 0.01; ***, p < 0.001.

To directly determine whether ATL1014 reduces the number of viral progenies, immunofluorescence analysis was performed on the lungs of SARS-CoV-2 infected hamsters to detect N protein. Lungs treated with ATL1014 exhibited significantly reduced viral antigens in pneumocytes and alveolar epithelium (Figure 5F). Correspondingly, the reduction in N protein^+^ signals correlated to the reduced levels of S protein, as shown by immunohistochemical analysis (Figure 5G). These results indicate that ATL1014 effectively reduces viral replication of SARS-CoV-2 *in vivo*.

### ATL1038 modulates gene expression to enhance autophagic activity and suppress SARS-CoV-2 replication

The pathogenesis of COVID-19 is associated with lung damages during acute illness, which involves pneumonia and, in severe cases, ARDS, and sepsis.^54^ Given the known physiological function of the ALS in inflammatory pathways,^55^ we tested whether ATL-bound p62 affects the gene expression involved in viral replication. SARS-CoV-2-infected Vero E6 cells were treated with ATL1038 and subjected to RNA-seq analyses (Figures 1A and S6A-S6B). Principal component analysis (PCA) revealed that the mRNA profiles of ATL1038-treated cells formed distinct clusters compared to that of vehicle-treated and uninfected groups (Figure 6B). The mRNA levels of 5,496 DEGs (differentially expressed genes) were significantly altered by SARS-CoV-2 infection (2,840 upregulated and 2,656 downregulated). ATL1038 treatment led to 1,997 DEGs (961 upregulated and 1,036 downregulated genes), and comparison revealed 1,211 overlapping DEGs. To gain further insights into biological significance, 1,211 DEGs were subjected to gene ontology (GO) enrichment analysis. Among the genes linked to SARS-CoV-2 infection, enriched GO terms were associated with transcription by RNA polymerase II, which facilitates the viral replication cycle and cis-regulatory region DNA binding.^56^ Other significant pathways involved in the infection process included TNF-alpha signaling via NF-kB and TNF signaling pathway (Figure 6C). However, upon treatment with ATL1038, the enriched GO terms were associated to the host’s antiviral defense mechanisms, including defense response to virus, ferroptosis, negative regulation of viral process, and regulation of viral genome replication. This result implies that ATL1038 modulates key antiviral pathways and may serve as a potential therapeutic agent for reducing viral replication. Additionally, treatment with ATL1038 led to enrichment of several GO terms related to the activation of autophagy, including autolysosome, and secondary lysosome pathways. Notably, heatmap analysis revealed a significant upregulation of key autophagy-related genes, including LAMP2, PLEKHM1, and SQSTM1. LAMP2, a maker of late stage autophagosomes, and PLEKHM1, which is involved in the recruitment of autophagic machinery, were both markedly increased, suggesting an enhancement of autophagic flux (Figure 6D). These results suggest that ATL1038 exerts antiviral activities through two synergistic mechanisms, by inducing the lysosomal degradation of viruses and modulating the transcriptional regulation of viral replication

**Figure 6.**
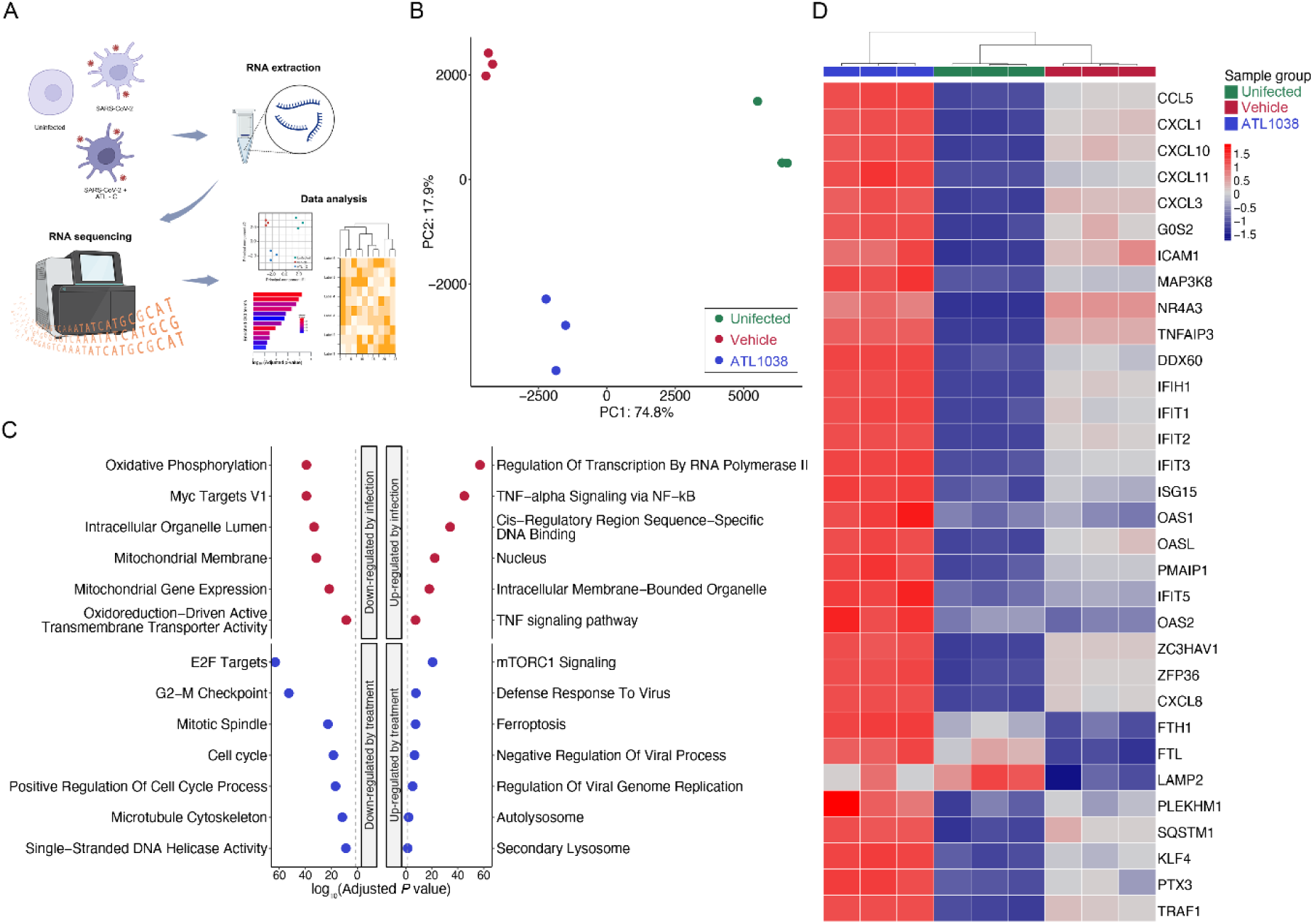
ATL1038 modulates gene expression to enhance autophagic activity and suppress SARS-CoV-2 replication. (A) Schematic of RNA seq analysis. (B) PCA plot of the 9 samples using gene expression data. (C) Dot plots showing significantly enriched GO terms for DEGs. Dotted line: adjusted *P = 0.05*. (D) Heatmap displaying expression pattern of the DEGs with an unsupervised hierarchical clustering of the samples.

## Discussion

Epidemics and pandemics are driven by the rapid changes of viral genetic information that enables them to evade host defense system. Traditional strategies in antiviral drug development have focused on the design of small molecules that target viral proteins, hopefully with high selectivity. However, the successful drugs often became no longer effective if mutations generate new variants. Recent studies explored various HTSs, in which drugs bind the components of host cells to achieve antiviral efficacy. If successful, HTS-based drugs may be effective for a broad range of viral strains and their variants. In this study, we explored the autophagic receptor p62 as a target for HTS-based antiviral. Our results showed that the autophagic activities of Vero E6 cells infected with SARS-CoV-2 were markedly elevated when treated with ATLs, as determined by the self-polymerization of p62 (Figure 1B) as well as synthesis and lipidation of LC3 (Figures S1A and S1B). These activities of ATLs were abolished in *p62^-/-^* MEFs (Figures 1D and 1E). Moreover, ATLs readily induced the targeting of viruses to autophagic membranes leading to their lysosomal co-degradation of SARS-CoV-2 (Figure 3C). These results demonstrate the HTS-based antiviral efficacy of ATLs that modulate p62 to target these viruses to autophagic membranes for lysosomal degradation.

One critical question remained to be addressed is the mechanism by which ATL-activated p62 selectively recognizes intracellular viruses. During the life-cycle of SARS-CoV-2, viral progenitors are assembled and matured within membranes derived from the ER or Golgi body.^57^ Following maturation, viruses encapsulated by SMVs are released from the host cells through exocytosis. Consistently, TEM analyses showed that autophagosomes predominantly sequester free viral particles (Figure 3C). It is therefore conceivable that p62 recognizes a pathogen-associated molecular pattern (PAMP) on viral surface. Intriguingly, TEM analyses also revealed that a minor portion of viruses are sequestered by autophagosomes as a vesicle-encapsulated form (data not shown), suggesting that ATL-activated p62 may also recognize a PAMP on SMVs. It is known that p62 can be associated with Lys63-linked Ub chains assembled on autophagic cargoes such as damaged mitochondria.^58^ We therefore predict that a similar mechanism may underlie in autophagic targeting of viruses by p62, a hypothesis to be validated in follow-up studies. We were not able to address these questions because experiments with SARS-CoV2 were conducted in the Biosafety Level 3 (BSL3) lab which was limited in Korea to access during the COVID-19 pandemic.

It has been reported that the main protease NSP5 of SARS-CoV-2 can cleave p62, preventing its ability to facilitate the degradation of viral proteins such as the M protein.^59^ Another study found that SARS-CoV-2 subverts ER-phagy by hijacking ER-phagy receptors, such as FAM134B and ATL3, into p62 condensates, leading to increased viral replication by promoting DMV formation.^60^ Previous finding has shown that autophagy activators upregulate autophagy-related genes which leads to re-establishing autophagy flux and exerts antiviral effects.^61^ Based on this, we hypothesized that activating p62 could overcome these virus-induced impairments. Our result demonstrates that ATLs successfully promote the recruitment of SARS-CoV-2 to autophagic membranes and restore lysosomal degradation. This approach highlights the potential of p62 activation as a novel strategy to counteract the viral escape mechanisms and enhance the degradation of viral particles via ALS.

It is known that intracellular SARS-CoV-2 induces autophagosome formation but subsequently impairs their maturation and fusion with lysosomes.^62–64^ As a part of its manipulation strategy to exploit host cellular machinery, the virus restricts autophagy-dependent protein degradation to prevent destruction of viral particles and utilizes autophagy-related lipid resources for viral production.^65^ It has been shown that ORF3a inhibits homotypic fusion and protein sorting (HOPS) complex from interacting with the SNARE protein which leads to failure of the formation of degradative autolysosomes.^63^ Similarly, another study demonstrated that ORF3a interacts with VPS39 disrupting the binding of HOPS to RAB7, thus inhibiting autophagosome-lysosome fusion and promoting accumulation of unfused autophagosomes.^66^ In our study, when SARS-CoV-2 infected cells were treated with ATL1038, LAMP and PLEKHM1 were markedly upregulated (Figure 6D). These findings suggest that ATL1038 not only restores autophagic flux but also facilitates the efficient fusion of autophagosomes with lysosomes.

As a family of *Coronaviridae*, SARS-CoV-2 infects respiratory tracts causing pneumonia-associated symptoms, during which lung epithelial cells and mucosal immune cells act as primary responders by producing interferons (IFNs), proinflammatory cytokines, and chemokines.^67^ Viral inflammation induced by these mediators is required to be tightly regulated to prevent cytokine storms that can exert severe tissue damage when excessive.^5^ To date, there are no effective drugs that counteract the replication and pathology of SARS-CoV-2 in the lung. Our results show that oral administration of ATL1014 significantly counteracted the replication of SARS-CoV-2 in the lungs of transgenic mice overexpressing hACE2. The antiviral activity of ATL1014 positively correlated to the viability and body weights in this acute model of SARS-CoV-2 (Figures 4B and 4C). Consistently, ATL1014 counteracted the viral replication as well as inflammatory lesions and other morphological abnormalities in the lungs of hamsters infected intranasally with SARS-CoV-2 (Figures 5B-5E). To our knowledge, ATL1014 is the first small molecule compound that exhibits therapeutic efficacy in viral replication and inflammatory lesions in the lungs of animal models. Given that ATL1014 is an orally administrative compound, chemical modulation of p62 may provide a potential antiviral strategy. Notably, the antiviral activity of ATLs in cultured cells was already comparable to those of drugs available in the market, including remdesivir (Figures S4A-SAC). It should be noted that the ATLs used in this study are hit compounds whose activities and physicochemical properties can be improved through structure-activity relationship studies. Indeed, our docking simulation study show that these ATLs bind p62 ZZ domain through in part common molecular interactions and in part distinct mode of interactions (Figure S2A-S2D). Further investigations are needed to understand the molecular interactions of ATLs and to increase their affinity and selectivity to p62 ZZ domain.

## Methods

### Cell culture and viruses

Vero E6, and HEK293T cell lines were obtained from American Type Culture Collection (ATCC). Wild-type and *p62^-/-^* mouse embryonic fibroblast (MEF) were kind gifts from Keji Tanaka (Tokyo Metropolitan Research Institute) with Tetsuro Ishii’s permission. Cell lines used in this study were maintained in Dulbecco’s Modified Eagle’s Medium (DMEM) supplemented with 10% FBS and 1% penicillin-streptomycin at 37[with 5% CO_2_. SARS-CoV-2 (NCCP No. 43326) was obtained from the National Culture Collection for Pathogens (Korea Disease Control and Prevention Agency), and a cell-adapted virus was generated by consecutively cultivating it in Vero E6 cells more than five times.

### Animal experiments

The mice experiments were approved by the Texas Biomedical Research Institute IACUC. Five-week-old female K18 hACE2 transgenic mice were purchased from The Jackson Laboratory. All animals were kept in the animal facility at Texas Biomedical Research Institute under specific pathogen-free conditions. Sixteen K18 hACE2 transgenic mice were randomly assigned to four mice per cage. Viral infection was performed through intranasally after gas anesthesia by isoflurane with reporter-expressing rSARS-CoV-2 at 1 x 10^4^ PFU. The mice were pre-treated with vehicle or ATL1014 a day before infection, followed by daily injection for 5 days. To examine the expression of Nluc, mice were anesthetized and the level of Nluc expression was measured via IVIS, and the expression of mCherry, pathological signs, and viral load were measured from lungs collected at 2 DPI.

The hamster experiments were approved by the Agency for Defense Development (ADD-IACUC-20-12). The infection experiments were performed at an approved biosafety level 3 facility. Five-week-old male Golden Syrian hamsters (Envigo, Indianapolis, IN, USA) were purchased from Raon Bio (Gyeonggi-do, South Korea). The hamsters were acclimatized for 7 days before the experiments. Fifteen hamsters were randomly assigned to three groups (n=5 hamsters per group) of which two groups were inoculated intranasally with SARS-CoV-2 (5 x 10^2^ PFU). The hamsters were pre-treated with vehicle or ATL1014 2 h before inoculation, followed by daily injection for 4 days. To examine pathological signs and viral load, lungs were collected 5 days after infection.

## Method details

### Synthesis of ATLs

ATL1014, 1024 and 1038 were used in this study along with the previously reported ATLs, ATL1002 and 1021, as positive control.

**Scheme 1.** Synthesis of **ATB1014**

^1^H NMR spectra were recorded on Bruker Avance III 400 MHz and Bruker Fourier 300 MHz and TMS was used as an internal standard.

LCMS was taken on a quadrupole Mass Spectrometer on Agilent 1260HPLC and 6120MSD (Column: C18 (50 × 4.6 mm, 5 μm) operating in ES (+) or (-) ionization mode; T = 30 °C; flow rate = 1.5 mL/min; detected wavelength: 220 nm.

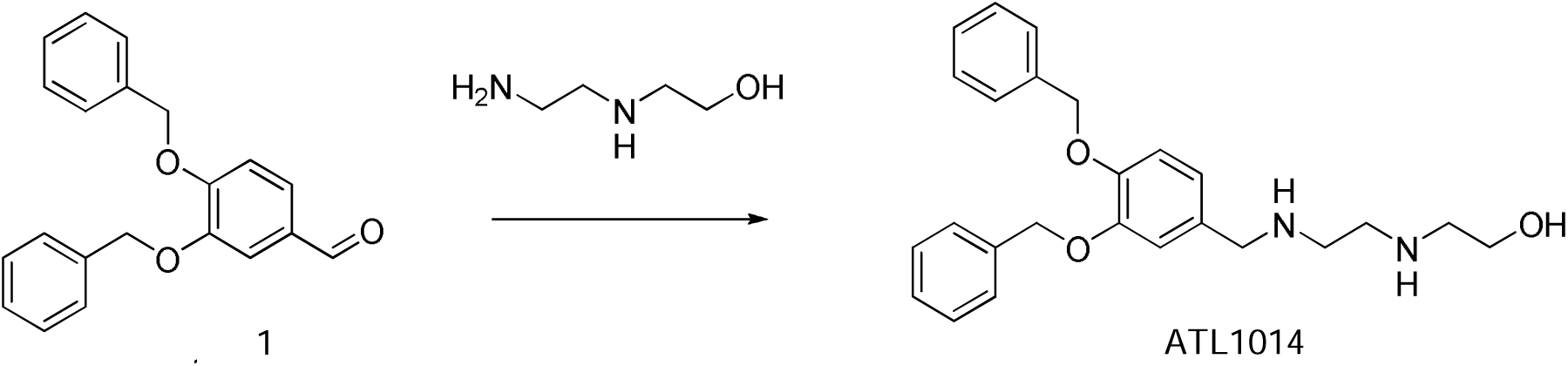

To a solution of **compound 1** (25.0 g, 78.5 mmol) in MeOH (250 mL) was added 2-(2-aminoethylamino)ethanol (9.00 g, 86.3 mmol). The mixture was **s**tirred overnight at 65 ^0^C. NaBH_4_ (3.50 g, 53.8 mmol) was added at 0 ^0^C. The reaction mixture was stirred 1 hour at rt. The above solution was poured into water. The solution was extracted with EtOAc (100 mL x 3). The combined organic layers were washed with brine, dried over Na_2_SO_4_ and concentrated. The crude was purified by prep-HPLC. The preparation solution was adjusted to pH 8-9 with solid NaHCO3 and extracted with EtOAc (100 mL x 6). The combined organic layers were dried over Na_2_SO_4_ and concentrated to give **ATL1014** (14.0 g) as yellow oil, yield: 43.8%.

^1^HNMR (CD_3_OD, 400 MHz): δ 7.44-7.48 (m, 4 H), 7.30-7.39 (m, 6 H), 7.11-7.12 (m, 1 H), 7.01-7.03 (m, 1 H), 6.91-6.93 (m, 1 H), 5.14-5.16 (m, 4 H), 3.79 (s, 2 H), 3.67-3.70 (m, 2 H), 2.86-2.89 (m, 2 H), 2.79-2.83 (m, 4 H).

LCMS [mobile phase: from 95% water (0.1% TFA) and 5% CH_3_CN to 40% water (0.1% TFA) and 60% CH_3_CN in 6.0 min, finally under these conditions for 0.5 min.] purity is >95%, Rt = 2.305 min; Mass Calcd.: 406; MS Found: 407 [MS+1].

**Scheme 2.** Synthesis of **ATL1024**

^1^H NMR spectra were recorded on Bruker Avance III 400 MHz and Bruker Fourier 300 MHz and TMS was used as an internal standard.

LCMS was taken on a quadrupole Mass Spectrometer on Agilent 1260HPLC and 6120MSD (Column: C18 (50 × 4.6 mm, 5 μm) operating in ES (+) or (-) ionization mode; T = 30 °C; flow rate = 1.5 mL/min; detected wavelength: 220 nm.

### 2.1 Synthesis of compound 3

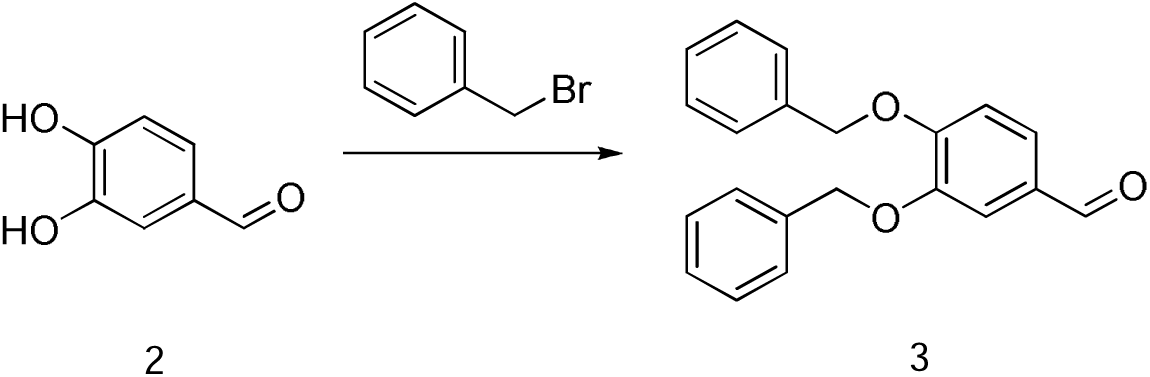

To a solution of **compound 2** (50 g, 0.37 mol) in ACN (750 mL) were added K_2_CO_3_ (200 g, 1.45 mol) and BnBr (137.5 g, 0.80 mol). The mixture was stirred at 85 °C for 16 hrs. The reaction was filtered and concentrated. The residue was added PE/EA (20:1, 50 mL) and stirred 1 h, filtered to afford **compound 3** (25 g) as white solid.

### 2.2 Synthesis of compound 4

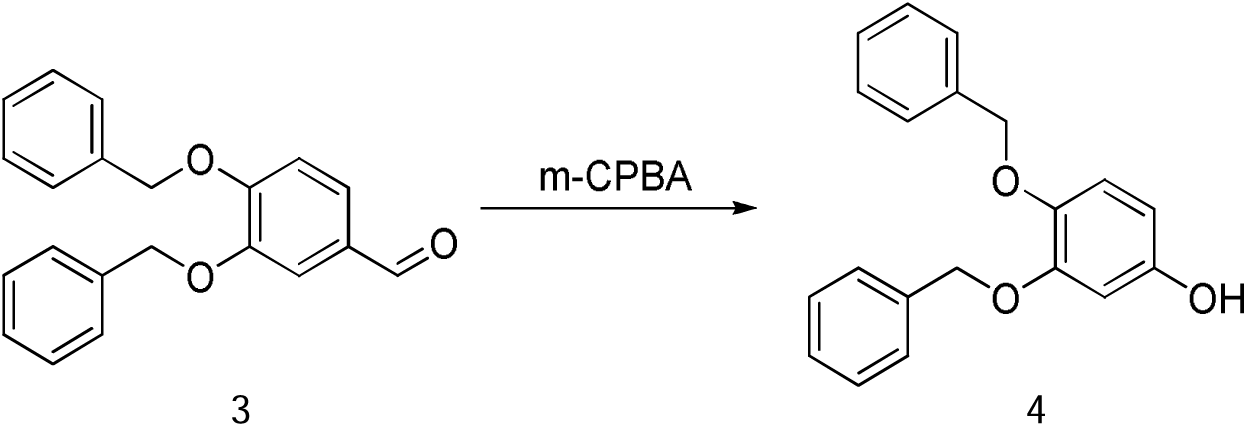

To a solution of **compound 3** (120 g, 377 mmol) in DCM (1000 mL) was added m-CPBA (129.6 g, 754 mmol). The mixture was stirred at room temperature for 16 hrs. Then the reaction was filtered. The filtration was concentrated. The residue was purified by silica gel, eluted with EA/PE (1:15∼1:5) to afford **compound 4** (62 g) as an off-white solid.

### 2.3 Synthesis of compound 5

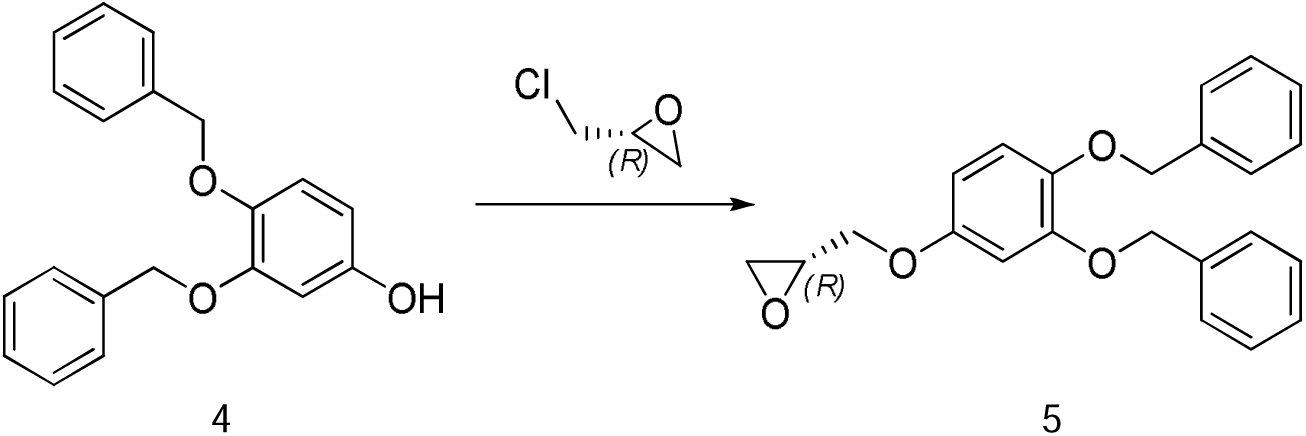

To a solution of **compound 4** (62 g, 202 mmol) in EtOH (1000 mL) were added water (40 mL) and KOH (18.8 g, 472 mmol). Then (R)-2-(chloromethyl)oxirane (55.8 g, 606 mmol) was added to the reaction. The resulting mixture was stirred at room temperature for 16 hrs. Then the reaction was quenched by addition water (1800 mL), extracted with EA (600 mL X 3). The organic layer was washed with brine, dried over Na_2_SO_4_, filtered and concentrated. The residue was purified by silica gel, eluted with EA/PE (1:15∼1:10) to afford **compound 5** (32 g) as a white solid.

### 2.4 Synthesis of compound 5

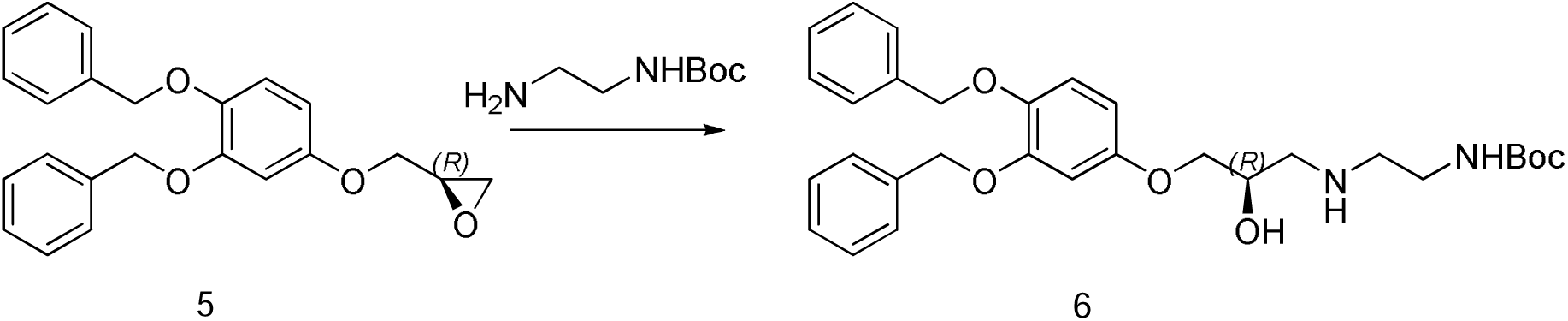

To a solution of **compound 5** (500 mg, 1.38 mmol) in MeOH (10 mL) were added tert-butyl 2-aminoethylcarbamate (442 mg, 2.76 mmol). The mixture was stirred at 50 °C for 10 hrs. Then the reaction was concentrated to afford **compound 6** (0.6 g) as a yellow solid without further purification.

### 2.4 Synthesis of compound 7

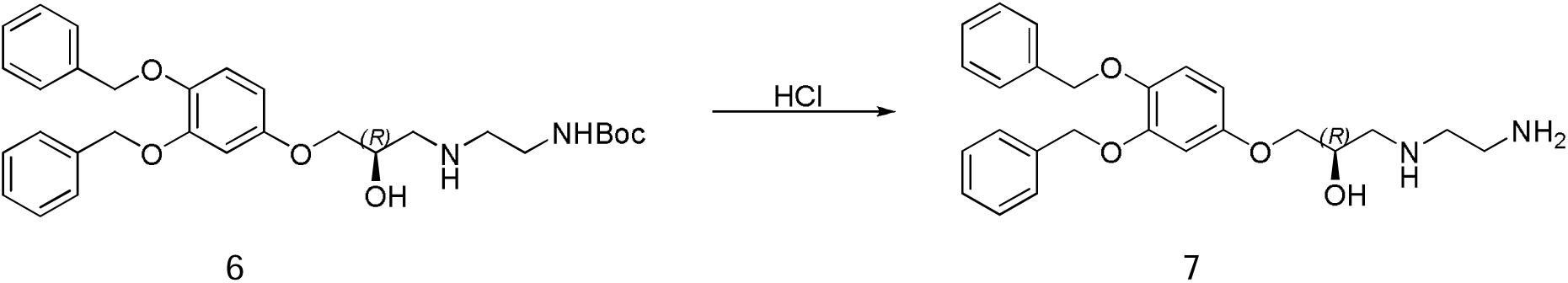

To a solution of **compound 6** (600 mg, 1.15 mmol) in MeOH (3 mL) was added MeOH/HCl (2 mL, 3N). The mixture was stirred at room temperature for 4 hrs. Then the reaction was concentrated to afford crude **compound 7** (400 mg) as a off-white solid, which was used directly for next step.

### 2.5 Synthesis of ATL1024

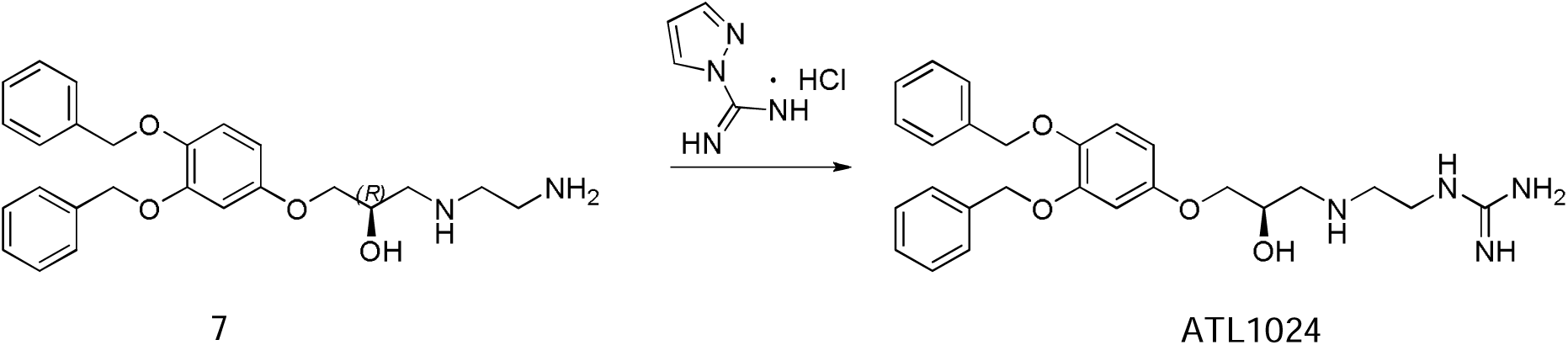

To a solution of **compound 7** (400 g, 0.95 mmol) in DMF (3 mL) were added 1H-Pyrazole-1-carboxamidine hydrochloride (553 mg, 3.8 mmol) and DIEA (0.3 g, 2.4 mmol). The mixture was stirred at 30 °C for 12 hrs. Then the solution was purified by pre-HPLC to afford **ATL1024** (22 mg) as a white solid.

**Scheme 3.** Synthesis of **ATL1038**

^1^H NMR spectra were recorded on Bruker Avance III 400 MHz and Bruker Fourier 300 MHz and TMS was used as an internal standard.

LCMS was taken on a quadrupole Mass Spectrometer on Agilent 1260HPLC and 6120MSD (Column: C18 (50 × 4.6 mm, 5 μm) operating in ES (+) or (-) ionization mode; T = 30 °C; flow rate = 1.5 mL/min; detected wavelength: 220 nm, 254nm.

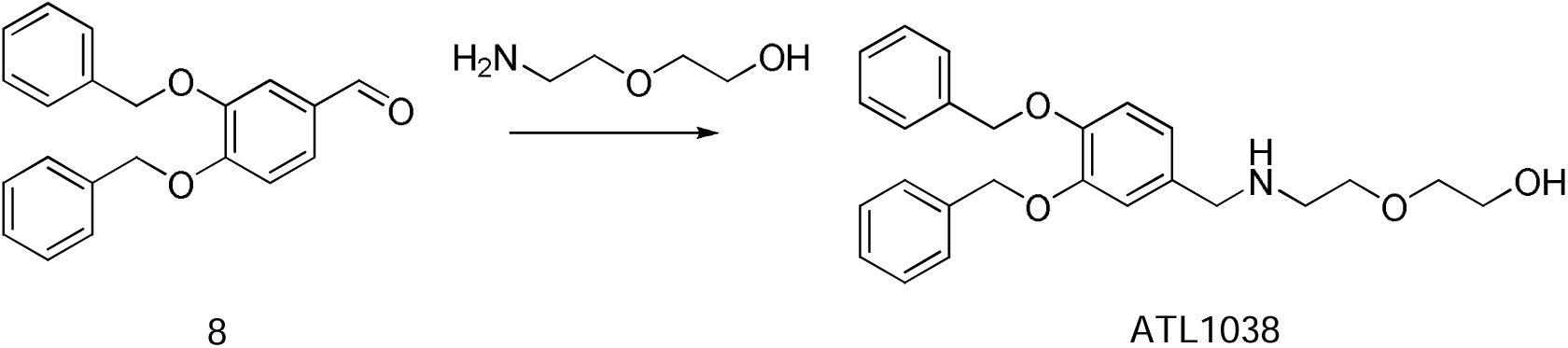

A mixture of **compound 8** (20.0 g, 62.8 mmol, 1.00 eq) added in MeOH (500 mL) and tetrahydrofuran (200 mL), Then added 2-(2-aminoethoxy)ethanol (13.2 g, 126 mmol, 2.00 eq) and sodium triacetoxyborohydrid (40.0 g, 189 mmol, 3.00 eq) at room temperature. The mixture reaction was stirred overnight at room temperature. The reaction mixture was quenched by the addition of saturated aqueous NH_4_Cl (200 mL). The aqueous layer was extracted with EA (200 mL), and concentrated. The crude was added PE (100 mL) and stirred for 1h, then filtered to give **ATL1038** (10.0 g, 39.1%) as a white solid. (TLC: DCM/MeOH=15/1, R_f_=0.5).

^1^HNMR (CDCl_3_, 400 MHz): δ 9.57 (s, 1H), 7.48 (d, *J* = 7.2 Hz, 2H), 7.27-7.41 (m, 9H), 6.99-7.02 (m, 1H), 6.84 (d, *J* = 8.4 Hz, 1H), 5.25 (s, 2H), 5.09 (s, 2H), 4.06(s, 1H), 3.65-3.70 (m, 4H), 3.46-3.48 (m, 2H), 2.87 (t, *J* = 4.8 Hz, 2H)

LCMS [mobile phase: from 90% water (0.05% TFA) and 10% CH_3_CN to 5% water (0.1% TFA) and 95% CH_3_CN in 6.0 min, finally under these conditions for 0.5 min.] purity is >98.6% (254 nm), Rt = 2.587 min; Mass Calcd.:407; MS Found: 408 [MS+1]. NMR and MS data could be found in the supplemental information.

### Docking simulation assay

For the docking simulation study, compounds were generated and optimized in Cresset Flare 24 software. Available crystal structure of p62 ZZ domain (PDB ID: 6MIU) was downloaded from Protein Data Bank (https://www.rcsb.org). Protein preparation was carried out in Cresset module Flare software. Hydrogen’s and 3D protonation were carried out on the target protein and minimized for the active site residues. Docking experiments were performed by using Cresset Flare software in accurate mode and default settings.

### *In vivo* p62 oligomerization assay

HEK 293T cells were treated with ATLs for 4 and 12 hours before harvesting. The harvested cells were then lysed using lysis buffer. Following centrifugation, the supernatants were collected, and protein concentrations were determined using a BCA protein assay kit. Equal amounts of protein from each lysate were mixed with non-reducing 4X LDS sample buffer and boiled. The samples were then separated using SDS-PAGE and oligomerized p62 was subsequently detected through immunoblotting analysis.

### Virus infection and small molecule ligands treatment

For infection, 0.01 MOI of SARS-CoV-2 was added to monolayer of Vero E6 cells with a confluence of 80∼90%. After incubation for 2 h at 37[, cells were rinsed with DPBS and then growth media containing the desired concentrations of small molecule ligands were added. The same concentration of DMSO was treated in negative control. After cultivation for the desired time at 37[, cells and supernatant were harvested respectively and analyzed for the purpose of each experiment.

### Dose-response curve analysis (DRC) by immunofluorescence

Ten-point DRCs were generated for each drug. Vero cells were seeded at 1.2 × 10^4^ cells per well in DMEM, supplemented with 2% FBS and 1X antibiotic-antimycotic solution (Gibco) in black, 384-well (Greiner Bio-One), 24 h prior to the experiment. Ten-point DRCs were generated, with compound concentrations ranging from 0.05–50 μM. For viral infection, plates were transferred into the BSL-3 containment facility and SARS-CoV-2 was added at a multiplicity of infection (MOI) of 0.0125. The cells were fixed at 24 hpi with 4% PFA and analyzed by immunofluorescence. The acquired images were analyzed using in-house software to quantify cell numbers and infection ratios, and antiviral activity was normalized to positive and negative controls in each assay plate. DRCs were fitted by sigmoidal dose response models, with the following equation: Y = Bottom + (Top Bottom) / (1 + (IC_50_ / X) Hillslope), using XLfit 4 Software. IC_50_ values were calculated from the normalized activity dataset-fitted curves. All IC_50_ and CC_50_ values were measured in duplicate, and the quality of each assay was controlled by Z’-factor and the coefficient of variation in percent (% CV).

### Plaque forming assay and plaque reduction assay

Plaque assay and plaque reduction assay were conducted as previously described ^68^. Briefly, SARS-CoV-2, which had been diluted in DMEM, was cultured in monolayer Vero E6 cells on a 24-well plate for 2 h at 37[. Cells were washed with DPBS and, for plaque assay, overlaid with 0.8% methylcellulose-DMEM mixture, containing 2% FBS. For plaque reduction assay, overlay media composed of 0.8% methylcellulose, DMEM, 2% FBS and the required concentration of small molecule ligands were added to cells. After 3 days of incubation at 37[, the plaques of SARS-CoV-2 were stained with rabbit anti-SARS-CoV-2 N protein antibody (AbFrontier) followed by being detected with alkaline phosphate-conjugated goat anti-rabbit IgG antibody (Invitrogen, 31340). The plaques were visualized and counted by incubation with nitro blue tetrazolium/5-bromo-4-chloro-3-indolyl-phosphate (NBT/BCIP) solution.

### Electron microscope analysis

For transmission electron microscopy, Vero E6 cells infected with SARS-CoV-2 were trypsinized and pelleted by centrifugation. Avoiding suspension, pellets were fixed in 2.5% glutaraldehyde in 0.1 M phosphate buffer overnight at 4[. Fixed pellets were embedded in epon resin and 55 nm sections were cut using ultramicrotome. Ultrathin sections were stained with uranyl acetate and lead citrate. Cell sections were examined using the 80 kV transmission electron microscope JEM-1400 (JEOL) at Electron Microscopy Center, Seoul National University Hospital Biomedical Research Institute.

### Immunoblotting

Cell pellets were washed with PBS and lysed in RIPA buffer supplemented with protease inhibitor cocktail and phosphatase inhibitor cocktail. Protein concentrations were quantified using the Pierce BCA Protein Assay Kit according to the manufacturer’s instructions. Protein supernatants were lysed in 5x Laemmli sample buffer. Whole-cell lysates were separated by SDS-PAGE and transferred onto polyvinylidene difluoride (PVDF) membranes. The membranes were blocked with 5% skim milk in 1X PBS-T buffer (PBS with 0.1% Tween-20) for 1 h at room temperature and incubated overnight with primary antibodies, followed by incubation with host-specific HRP-conjugated secondary antibodies. For signal detection, a mixture of ECL solution was applied onto the membrane and captured using X-ray films.

### Immunocytochemistry

Cells were cultured on coverslips coated with poly L-lysine solution in 24-well plates. The cells were fixed with 4% PFA in PBS for 15 mins at room temperature and washed three times with PBS. The cells were permeabilized with 0.5% Triton X-100 in PBS and washed three times with PBS. The cells were blocked with 2% BSA in PBS for 1 h at room temperature and incubated overnight with primary antibodies in 2% BSA in PBS at 4[. After incubation, the cells were washed three times with PBS and incubated with alexa fluor-conjugated secondary antibodies in a blocking buffer for 1 h at room temperature. Subsequently, the coverslips were mounted on glass slides using a DAPI-containing mounting medium. Confocal images were acquired by laser scanning confocal microscope 510 Meta (Zeiss) and analyzed by Zeiss LSM Image Browser (ver. 4.2.0.121).

### Histological examination

The left lungs of each hamster were fixed in 4% paraformaldehyde phosphate buffer solution and processed for paraffin embedding. The paraffin blocks were cut into 5-μm-thick sections. Two sections from each sample were stained using a standard hematoxylin and eosin (H&E) procedure. Histopathological changes were determined by a certified pathologist.

### Immunohistochemistry

Lung sections were deparaffinized and rehydrated following antigen retrieval with citrate buffer (pH 6.0) at 98[and washed three times with PBS. For quenching endogenous peroxidase activity, tissue sections were agitated in 3% H_2_O_2_ diluted in PBS for 15 mins at room temperature and washed with distilled water three times. Tissue sections were blocked in blocking solution (2% NGS, 1% BSA, 0.1% triton X-100, and 0.05% Tween-20 in PBS) for 1 h at room temperature. After blocking, tissue sections were incubated with diluted primary antibodies in a humidified chamber overnight at 4[. After incubation, tissue sections were washed with PBS and incubated in HRP-conjugated secondary antibody for 1 h at room temperature. After washing, tissue sections were incubated with DAB substrate kit following the manufacturer’s instructions. Developed tissue sections were counter-stained with hematoxylin and mounted with mounting medium. Images were obtained using a digital slide scanner (Axio Scan, Zeiss).

### Transcriptome analysis

The libraries for RNA-seq were constructed using SMART-Seq ^®^ v4 Ultra ^®^ low input kit and were sequenced using Illumina to generate 101 bp paired-end reads. The resulting reads were aligned to the combined genome of Chlorocebus sabaeus (GCF_000409795.2, NCBI) and COVID-19 (ASM985889v3, NCBI), using the RSEM (version 1.3.3.). Read counts were transformed using the DESeq2 (version 1.28.1), and significant DEGs were identified by adjusted *P* < 0.05 and absolute value of log_2_ fold change < 1.2. To perform PCA, R function ‘prcomp’ was used with the gene expression values from RSEM. GO enrichment analysis was performed using the R package ‘enrichR’.

### Quantification and statistical analysis

All statistical analyses were performed using GraphPad Prism 5. Results are reported as mean ± SD. Significant differences between 2 groups were performed by the 2-tailed Student’s *t* test. Differences between groups were considered statistically significant for *P* < 0.05.

## Acknowledgments

We thank the laboratory members of Y.T.K. and AUTOTAC Bio Inc. for their critical discussion and comments during this study. The antiviral efficacy of ATLs in cultured cells were conducted at Institute Pasteur Korea during the COVID-19 pandemic through the support by the Korean government. This work was supported by the National Research Foundation of Korea (NRF) grants funded by the Ministry of Science and ICT (MSIP) (2020R1A5A1019023 to Y.T.K., 2021M3A9I2080489 to N.H.C., and 2021M3A9I2080490 to N.H.C.), and the Ministry of Education (2021R1A2B5B03002614 to Y.T.K.). This study was also supported from the Korea-US Collaborative Research Project funded by MSIP and Future Planning and Ministry of Health and Welfare of Korea (RS-2024-00467046 to N.H.C.).

## Author contributions

Conceptualization, T.H.B., G.E.L., U.P., K.W.S., N.H.C., and Y.T.K.; methodology, T.H.B., G.E.L., U.P., J.L., D.W.L., and K.W.S.; formal analysis, T.H.B., G.E.L., and K.W.S.; investigation, T.H.B., G.E.L., U.P., Y.J.S., S.H.G., C.H.Y., M.K., L.M., M.C., J.P., and K.W.S.; resources, M.C., J.P., N.H.C., and Y.T.K.; writing – original draft, K.W.S., G.E.L, J.L., and Y.T.K.; writing – review & editing, G.E.L, U.P., M. K., M.C., and Y.T.K.; visualization, K.W.S., and G.E.L.; supervision and project administration, Y.T.K. All authors contributed to the article and approved the submitted version.

## Data and materials availability

All data are available in the main text or the supplementary materials.

## Supplemental Information

Supplementary Figure S1. Chemical mimetics of Arg/N-degrons induce the lysosomal degradation of SARS-CoV-2 via p62-dependent autophagy

Supplementary Figure S2. Docking models of the small molecule ligands to p62 ZZ domain

Supplementary Figure S3. Cytotoxicity of ATLs

Supplementary Figure S4. Dose-response curve analysis of N protein for reference antivirals

Supplementary Figure S5. RNA-seq quality assessment

Supplementary Figure S1. Chemical mimetics of Arg/N-degrons induce the lysosomal degradation of SARS-CoV-2 via p62-dependent autophagy

(A) Immunoblotting analysis of autophagy flux upon SARS-CoV-2 infection. Vero E6 Cells infected with SARS-CoV-2 for 2 h were treated with ATL1038 for 12 h followed by combinational treatment with 200 nM bafilomycin A1 for 4h. (B) Immunofluorescence analysis of autophagy flux. Infected Vero E6 cells were treated with ATL1038 for 12 h at 5 μM followed by combinational treatment with 200 nM bafilomycin A1 for 4 h. (C) LC3 puncta formation assay. VeroE6 cells were infected with SARS-CoV-2 and incubated for 14 h, followed by immunostaining analysis. (D) Quantitation of (C).

Supplementary Figure S2. Docking models of the small molecule ligands to p62 ZZ domain

(A) The crystal structure of p62 ZZ domain in complex with Arg-Glu peptide (PDB ID: 6MIU). (B-D) The binding modes of ATL1014 (B), ATL1024 (C), and ATL1038 (D)

Supplementary Figure S3. Cytotoxicity analysis of ATLs

(A-C) Dose-response curve analysis for cytotoxicity of ATL1014 (A), ATL1024 (B), and ATL 1038 (C)

Supplementary Figure S4. Dose-response curve analysis of N protein for reference drugs

(A-C) Dose-response curve of chloroquine (A), remdesivir (B), and lopinavir (C). Reference drugs were added to Vero E6 cells prior to the infection with SARS-CoV-2 at a MOI of 0.0125. The immunofluorescence images were obtained to generate the dose-response curve at 24 hpi.

Supplementary Figure S5. RNA-seq quality assessment

A. Distribution of transcripts per millions mapped reads (TPM) from 9 samples. (B) Expression of viral ORFs in the samples were analysed for RNA-seq.

