## Supplemental Figures 1,2,3,4,5 for "Targeted degradation of SARS-CoV-2 via the autophagy-lysosome system using chemical mimetics of the N-degron pathway"

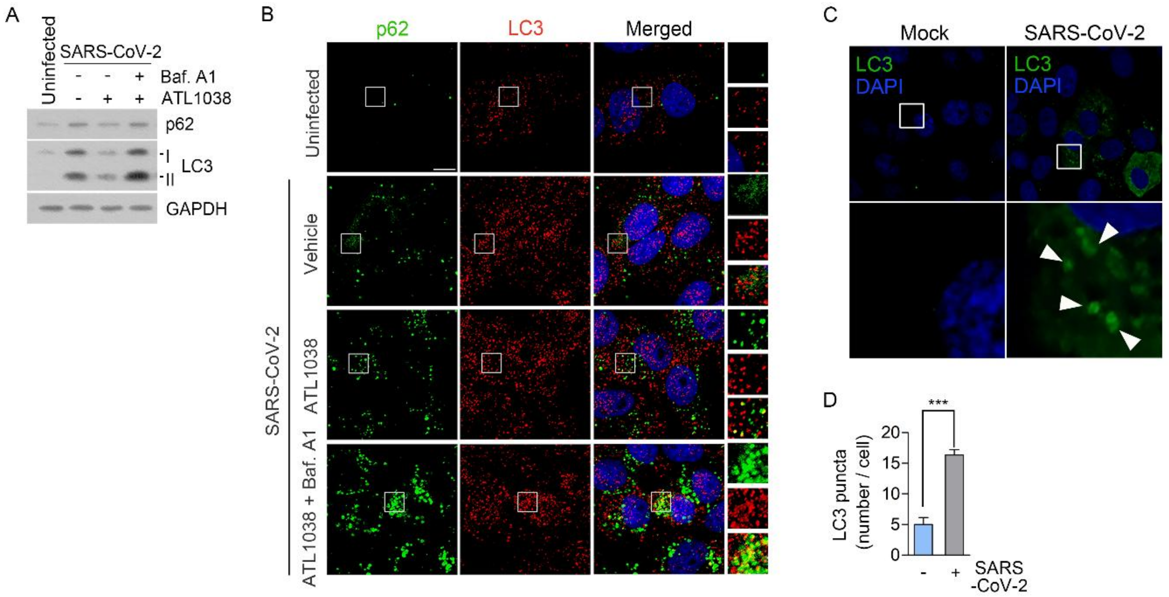


**Supplementary Figure S1. Chemical mimetics of Arg/N-degrons induce the lysosomal degradation of SARS-CoV-2 via p62-dependent autophagy**

(A) Immunoblotting analysis of autophagy flux upon SARS-CoV-2 infection. Vero E6 Cells infected with SARS-CoV-2 for 2 h were treated with ATL1038 for 12 h followed by combinational treatment with 200 nM bafilomycin A1 for 4h. (B) Immunofluorescence analysis of autophagy flux. Infected Vero E6 cells were treated with ATL1038 for 12 h at 5 μM followed by combinational treatment with 200 nM bafilomycin A1 for 4 h. (C) LC3 puncta formation assay. VeroE6 cells were infected with SARS-CoV-2 and incubated for 14 h, followed by immunostaining analysis. (D) Quantitation of (C).

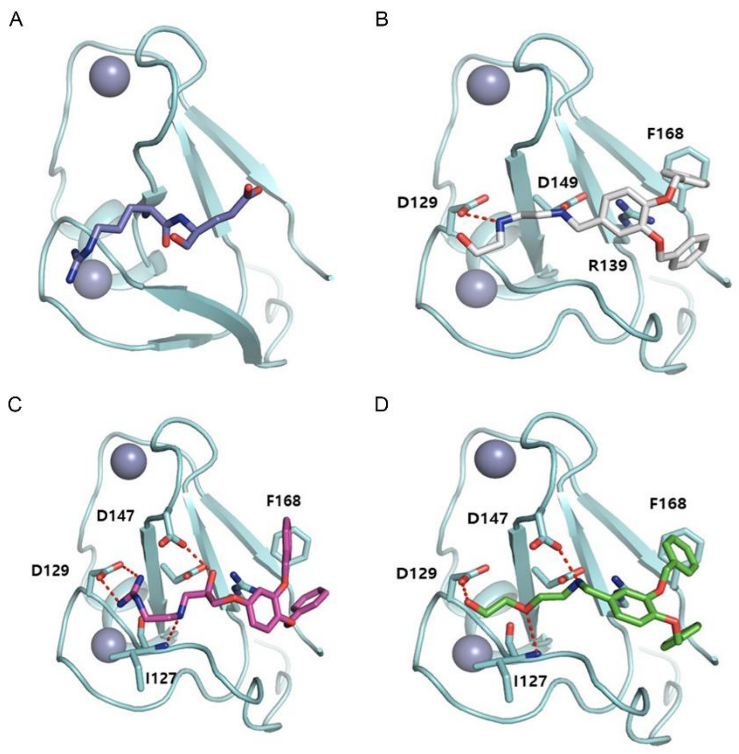


**Supplementary Figure S2. Docking models of the small molecule ligands to p62 ZZ domain**

(A) The crystal structure of p62 ZZ domain in complex with Arg-Glu peptide (PDB ID: 6MIU). (B-D) The binding modes of ATL1014 (B), ATL1024 (C), and ATL1038 (D)

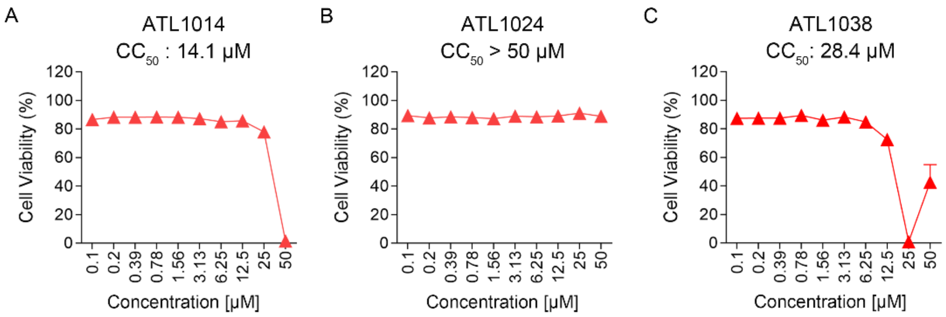


**Supplementary Figure S3. Cytotoxicity analysis of ATLs**

(A-C) Dose-response curve analysis for cytotoxicity of ATL1014 (A), ATL1024 (B), and ATL 1038 (C)

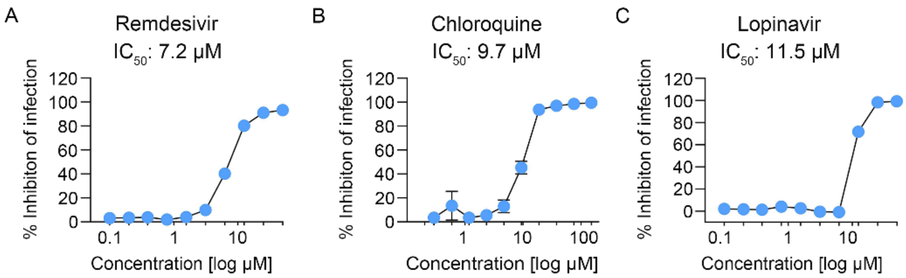


**Supplementary Figure S4. Dose-response curve analysis of N protein for reference drugs**

(A-C) Dose-response curve of chloroquine (A), remdesivir (B), and lopinavir (C). Reference drugs were added to Vero E6 cells prior to the infection with SARS-CoV-2 at a MOI of 0.0125. The immunofluorescence images were obtained to generate the dose-response curve at 24 hpi.

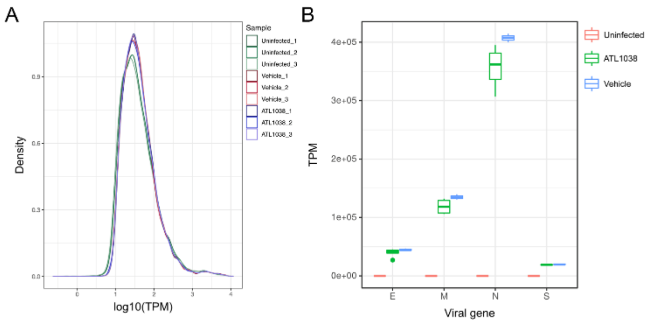


**Supplementary Figure S5. RNA-seq quality assessment**

(A) Distribution of transcripts per millions mapped reads (TPM) from 9 samples. (B) Expression of viral ORFs in the samples were analysed for RNA-seq.
